# ERGA-BGE Reference Genome of *Loboptera subterranea*: a cave adapted cockroach endemic to the Canary Islands

**DOI:** 10.1101/2025.02.06.636782

**Authors:** Heriberto López, Pedro Oromí, Xavier Belles, Guillem Ylla, Nuria Escudero, Rosa Fernández, Astrid Böhne, Marta Gut, Laura Aguilera, Tyler Alioto, Francisco Câmara Ferreira, Jèssica Gómez-Garrido, Fernando Cruz, Rita Monteiro

## Abstract

Species of the genus *Loboptera* from the Canary Islands are adapted to life in subterranean environments, except *Loboptera canariensis* and *Loboptera decipiens* which inhabit epigean, often influenced by human activities. The genome of *Loboptera subterranea* provides a valuable resource for understanding the genetic basis of how the Canarian *Loboptera* species evolved and thrived in subterranean environments.

A total of 14 contiguous chromosomal pseudomolecules were assembled from the genome sequence. This chromosome-level assembly encompasses 2.7 Gb, composed of 486 contigs and 90 scaffolds, with contig and scaffold N50 values of 13.1 Mb and 202.7 Mb, respectively.

## Introduction

*Loboptera subterranea* (Blattodea, Blattellidae) is one of the 13 species of cockroaches of the genus *Loboptera* found in the Canary Islands. All *Loboptera* species occurring in these islands, except for *Loboptera canariensis* and *Loboptera decipiens*, are endemics and have a variable degree of adaptation to subterranean life, especially relating to pigmentation and eye reduction (Martín & Oromí, 1987; Martín & Izquierdo, 1987; Izquierdo & Martín, 1990; Martín et al., 1999). *Loboptera troglobia* and *L. subterranea* are the most specialized species showing a remarkable depigmentation and complete anophthalmia, as well as other troglomorphic characters such as reduction of the number of ovarioles and compartments in the ootheca (Izquierdo et al., 1990). *Loboptera subterranea* has been found in caves of the North of Tenerife island, in some locations together with *L. troglobia* (Martín et al., 1999). Both are relevant species of the subterranean habitats of Tenerife and probably play a prominent role in this ecosystem.

The hypogean (= subterranean) species of Canarian *Lobopter*a have derived from an epigean (= surface) ancestor after having adapted to the subterranean environment by undergoing drastic morphological and physiological transformations (Izquierdo, 1997; Martín et al., 1999). The study of the genome of *L. subterranea* and its comparison with that of the epigean *L. canariensis* and *L. decipiens* may reveal to what extent evolutionary mechanisms operate to produce transformations that enable adaptation to live in the underground environment.

The epigean ancestor of *L. subterranea* is unknown, given that *L. canariensis* and *L. decipiens* are probably introduced and the only epigean species occurring on the Canary Islands. On the other hand, an unpublished study indicates that the hypogean species of Canarian *Loboptera* are more closely related to the epigean species of Morocco than to *L. canariensis* and *L. decipiens* (Izquierdo, 1997). *L subterranea* is one of seven completely eyeless hypogean species of the genus in the Canary Islands, which makes its genome interesting considering that the other five endemic hypogean species have reduced but still present eyes.

The generation of this reference resource was coordinated by the European Reference Genome Atlas (ERGA) initiative’s Biodiversity Genomics Europe (BGE) project, supporting ERGA’s aims of promoting transnational cooperation to promote advances in the application of genomics technologies to protect and restore biodiversity (Mazzoni et al., 2023).

## Materials & Methods

ERGA’s sequencing strategy includes Oxford Nanopore Technology (ONT) and/or Pacific Biosciences (PacBio) for long-read sequencing, along with Hi-C sequencing for chromosomal architecture, Illumina Paired-End (PE) for polishing (i.e. recommended for ONT-only assemblies), and RNA sequencing for transcriptomic profiling, to facilitate genome assembly and annotation.

### Sample and Sampling Information

On September 26, 2023, one adult female and one adult male individual of *Loboptera subterranea* were sampled by Pedro Oromí and Heriberto López. Identification was based on Martín et al. (1999) and performed by Pedro Oromí and Heriberto López. The species identification through COI barcoding was done at the National Museum of Natural Sciences (CSIC). The specimen was directly searched and captured by hand in Icod de los Vinos (Tenerife) at Sobrado Superior Cave. Sampling was performed under permission Ref. 2022/9790 and Ref. 2023-00957 issued by the Government of The Canary Islands and the Cabildo of Tenerife respectively. The thorax, head, and abdomen of the specimens were snap-frozen immediately after harvesting and stored in liquid nitrogen until DNA extraction.

### Vouchering information

Physical reference materials for a proxy specimen have been deposited in the Museo Nacional de Ciencias Naturales (MNCN, CSIC) https://mncn.csic.es/en under the accession number MNCN_Ent 375554.

Frozen reference tissue material from leg is available from both individuals at the Biobank of the Museo Nacional de Ciencias Naturales under the voucher IDs MNCN-ADN-151766 (adult female) and MNCN-ADN-151767 (adult male).

### Data Availability

*Loboptera subterranea* and the related genomic study were assigned to Tree of Life ID (ToLID) ‘ibLobSubt1’ and all sample, sequence, and assembly information are available under the umbrella BioProject PRJEB77747. The sample information is available at the following BioSample accessions: SAMEA114562287 and SAMEA114562292. The genome assembly is accessible from ENA under accession number GCA_964278025.1 Sequencing data produced as part of this project are available from ENA at the following accessions: ERX13167079, ERX12745425 and ERX13167080. Documentation related to the genome assembly and curation can be found in the ERGA Assembly Report (EAR) document available at https://github.com/ERGA-consortium/EARs/blob/main/Assembly_Reports/Loboptera_subterranea/ibLobSubt1/ibLobSubt1_EAR.pdf. Further details and data about the project are hosted on the ERGA portal at https://portal.erga-biodiversity.eu/data_portal/3078426.

### Genetic Information

The estimated genome size, based on ancestral taxa, is 1.49 Gb. This is a diploid genome with an estimated haploid number based on ancestral taxa of 17 chromosomes (2n=34). Information for this species was retrieved from Genomes on a Tree (Challis et al., 2023).

### DNA/RNA processing

DNA was extracted from the abdomen of one specimen using the Blood & Cell Culture DNA Mini Kit (Qiagen) following the manufacturer’s instructions. DNA quantification was performed using a Qubit dsDNA BR Assay Kit (Thermo Fisher Scientific), and DNA integrity was assessed using a Genomic DNA 165 Kb Kit (Agilent) on the Femto Pulse system (Agilent). The DNA was stored at +4oC until used.

RNA was extracted using the RNeasy Mini Kit (Qiagen) according to the manufacturer’s instructions. RNA was extracted from two different specimen parts: the abdomen and thorax. RNA quantification was performed using the Qubit RNA HS kit and RNA integrity was assessed using a Bioanalyzer 2100 system (Agilent) RNA 6000 Pico Kit (Agilent). Equimolar amounts of RNA were pooled for the library preparation and stored at −80°C until used.

### Library Preparation and Sequencing

For long-read whole genome sequencing, a library was prepared using the SQK-LSK114 Kit (Oxford Nanopore Technologies, ONT) and was sequenced on a PromethION 24 A Series instrument (ONT). For short-read whole genome sequencing (WGS), a library was prepared using the KAPA Hyper Prep Kit (Roche). A Hi-C library was prepared from the abdomen and thorax using the Dovetail Omni-C Kit (Cantata Bio), followed by the KAPA Hyper Prep Kit for Illumina sequencing (Roche). The RNA library was prepared from the pooled sample using the KAPA mRNA Hyper prep kit (Roche). The short-read libraries were sequenced on a NovaSeq 6000 instrument (Illumina). In total, 46X Oxford Nanopore, 38X Illumina WGS shotgun, and 41X HiC data were sequenced to generate the assembly.

### Genome Assembly Methods

The genome was assembled using the CNAG CLAWS pipeline (Gomez-Garrido, 2024). Briefly, reads were preprocessed for quality and length using Trim Galore v0.6.7 and Filtlong v0.2.1, and initial contigs were assembled using NextDenovo v2.5.0, followed by polishing of the assembled contigs using HyPo v1.0.3, removal of retained haplotigs using purge-dups v1.2.6 and scaffolding with YaHS v1.2a. Finally, assembled scaffolds were curated via manual inspection using Pretext v0.2.5 with the Rapid Curation Toolkit (https://gitlab.com/wtsi-grit/rapid-curation) to remove any false joins and incorporate any sequences not automatically scaffolded into their respective locations in the chromosomal pseudomolecules (or super-scaffolds). Finally, the mitochondrial genome was assembled as a single circular contig of 16,433 bp using the FOAM pipeline v0.5 (https://github.com/cnag-aat/FOAM) and included in the released assembly (GCA_963981305.1). Summary analysis of the released assembly was performed using the ERGA-BGE Genome Report ASM Galaxy workflow (https://workflowhub.eu/workflows/1103?version=3), incorporating tools such as BUSCO v5.5, Merqury v1.3, and others (see reference for the full list of tools).

## Results

### Genome Assembly

The genome assembly has a total length of 2,671,328,670 bp in 91 scaffolds including the mitogenome (Figures 1 and 2), with a GC content of 34.38%. It features a contig N50 of 13,138,595 bp (L50=60) and a scaffold N50 of 202,725,563 bp (L50=6). There are 396 gaps, totaling 79,200 kb in cumulative size. The single-copy gene content analysis using the arthropoda_odb10 database with BUSCO resulted in 99.0% completeness (97.2% single and 1.8% duplicated). 86.5 % of reads k-mers were present in the assembly and the assembly has a base accuracy Quality Value (QV) of 41.8 as calculated by Merqury.

**Figure 1.**
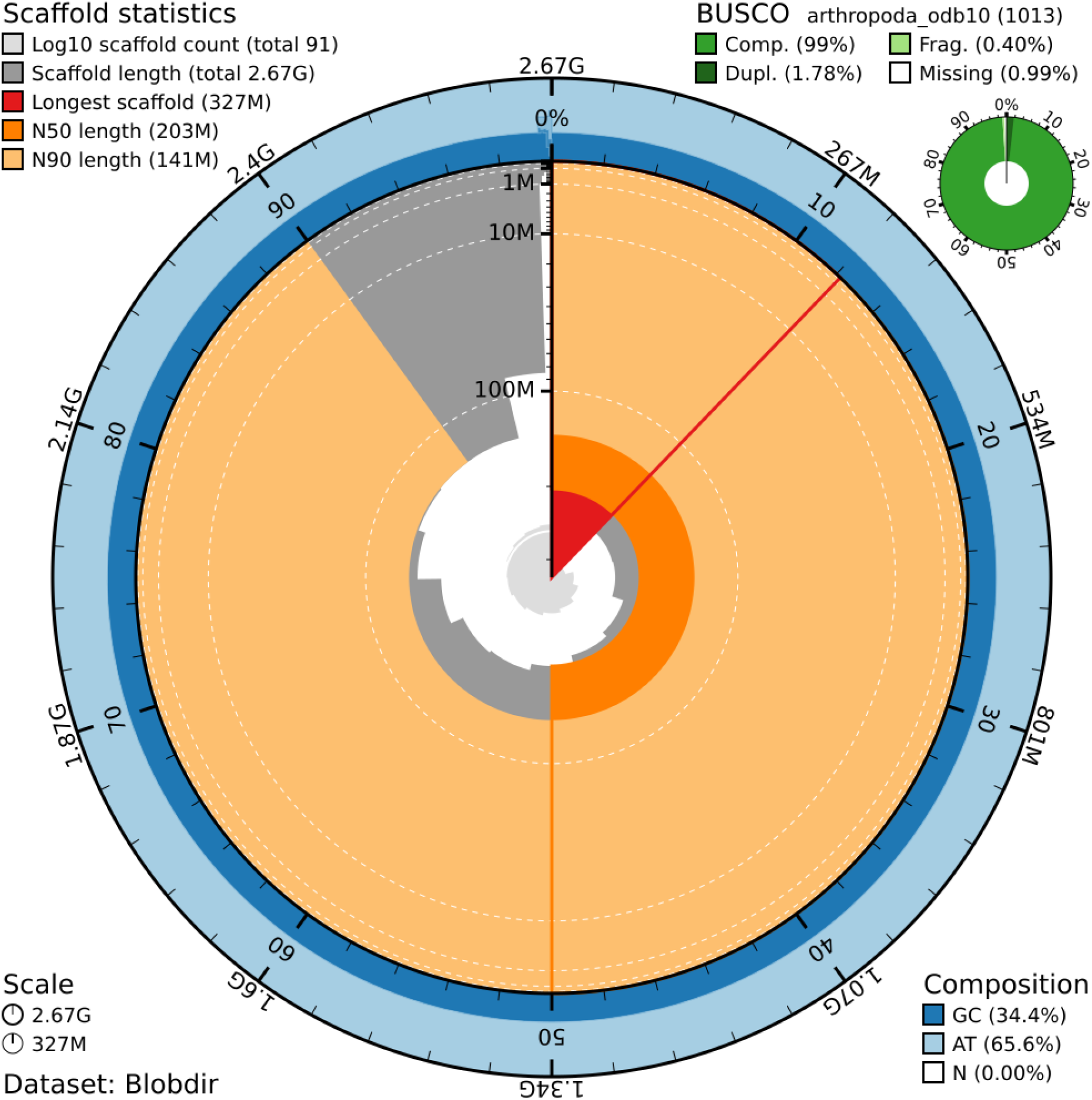
Snail plot summary of assembly statistics. The main plot is divided into 1,000 size-ordered bins around the circumference, with each bin representing 0.1% of the 2,671,328,670 bp assembly including the mitochondrial genome. The distribution of sequence lengths is shown in dark grey, with the plot radius scaled to the longest sequence present in the assembly 327 bp, shown in red). Orange and pale-orange arcs show the scaffold N50 and N90 sequence lengths (203 and 141 bp), respectively. The pale grey spiral shows the cumulative sequence count on a log-scale, with white scale lines showing successive orders of magnitude. The blue and pale-blue area around the outside of the plot shows the distribution of GC, AT, and N percentages in the same bins as the inner plot. A summary of complete, fragmented, duplicated, and missing BUSCO genes found in the assembled genome from the Arthropoda database (odb10) is shown on the top right.

**Figure 2.**
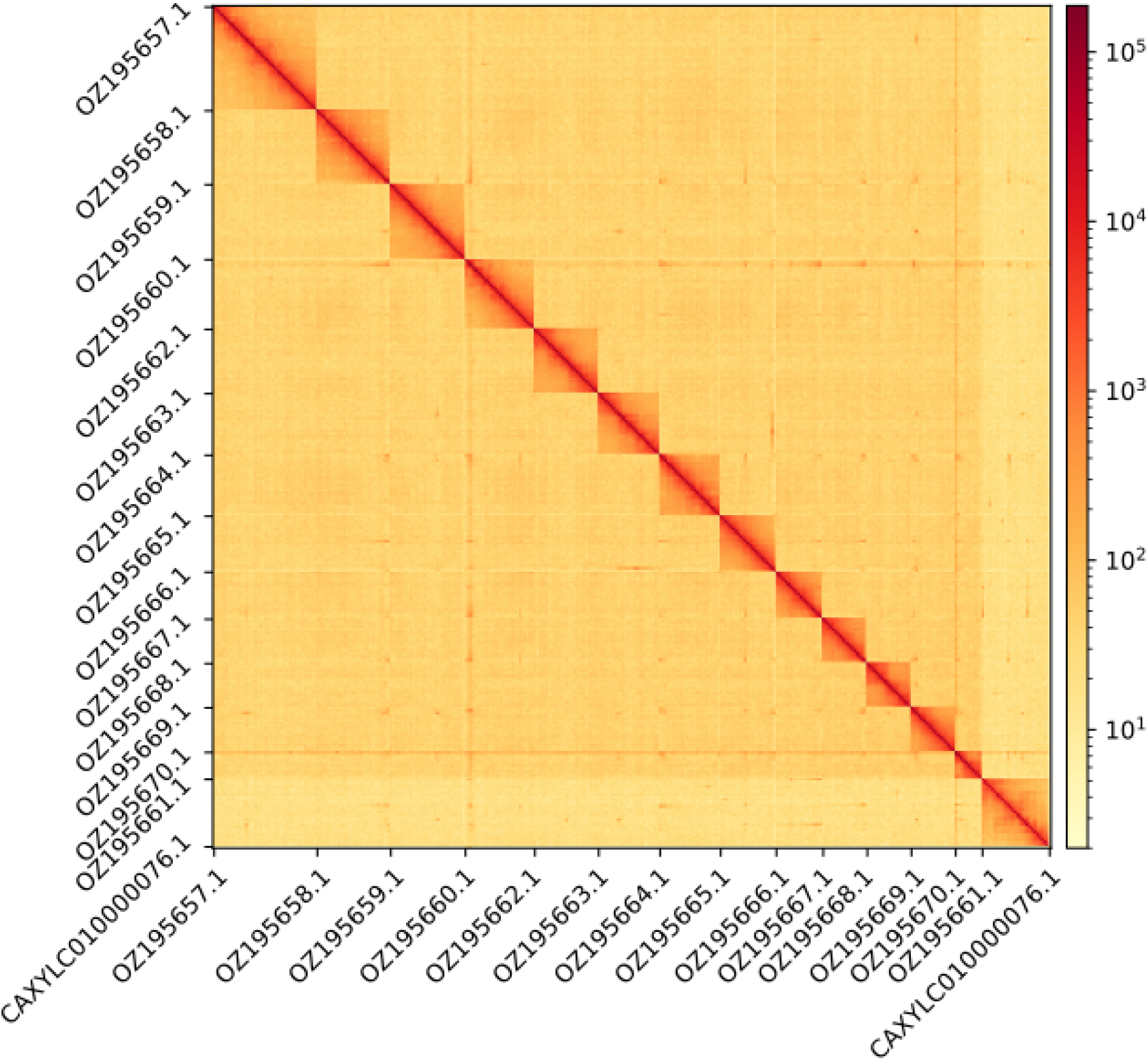
Hi-C contact map showing spatial interactions between regions of the genome. The diagonal corresponds to intra-chromosomal contacts, depicting chromosome boundaries. The frequency of contacts is shown on a logarithmic heatmap scale. Hi-C matrix bins were merged into a 150 kb bin size for plotting.

## Acknowledgements

We would like to express our gratitude to Nuria Macías, Daniel Febles, Antonio J. Pérez, David Lugo, Diego Patiño, Pablo Caloca and Francisco Mesa for their assistance in the field, and also to Esther Martín for managing the permits to lead us to access to Cueva del Sobrado.

We acknowledge the support of the Freiburg Galaxy Team: Saim Momin and Björn Grüning, Bioinformatics, University of Freiburg (Germany), funded by the German Federal Ministry of Education and Research BMBF grant 031 A538A de.NBI-RBC and the Ministry of Science, Research and the Arts Baden-Württemberg (MWK) within the framework of LIBIS/de.NBI Freiburg. We would like to acknowledge the assembly reviewer, Michael Paulini from The Wellcome Sanger Institute.

## Author Contributions

PO, HL collected the species, PO, HL identified the species, PO and HL sampled and preserved biological material and provided metadata, RM, AsB, NE, and RF provided support in sampling, shipping of biological material, metadata collection, and management, LA and MG extracted DNA, prepared libraries, and performed sequencing, FCF, JGG and FC performed genome assembly and curation under the supervision of TA, RM generated the analysis and report. All authors contributed to the writing, review, and editing of this genome note and read and approved the final version.

## Literature Cited

Challis R, Kumar S, Sotero-Caio C, Brown M & Blaxter M (2023) Genomes on a Tree (GoaT): A versatile, scalable search engine for genomic and sequencing project metadata across the eukaryotic tree of life [version 1; peer review: 2 approved]. Wellcome Open Research 8:24. 10.12688/wellcomeopenres.18658.1

Gomez-Garrido J (2024) CLAWS (CNAG’s Long-read Assembly Workflow in Snakemake) [Computer software]. 10.48546/WORKFLOWHUB.WORKFLOW.567.2

Izquierdo l (1997) Estrategias adaptativas al medio subterráneo de las especies del género Loboptera Brunner W. (Blattaria, Blattellidae) en las Islas Canarias. Tesis Doctoral. Universidad de La Laguna, Tenerife, 324 pp.

Izquierdo I & Martín JL (1990) Una nueva especie anoftalma de Loboptera Brunner W. en la isla de Tenerife (Islas Canarias) (Blattaria, Blattellidae). Fragmenta Entomologica, 22 (1): 19–25.

Izquierdo I, Oromí P & Belles X (1990) Number of ovarioles and degree of dependence with respect to the underground environment in the Canarian species of the genus Loboptera Brunner (Blattaria, Blattellidae). Mém. Biospéol., 17: 107–111.

Martín JL & Izquierdo I (1987) Dos nuevas formas hipogeas de Loboptera (Blattaria, Blattellidae) en la isla de El Hierro (Islas Canarias). Fragmenta Entomologica, 19 (2): 301–310.

Martín J L, Izquierdo I & Oromí P (1999) El género Loboptera en Canarias; descripción de cinco nuevas especies hipogeas (Blattaria: Blattellidae). Vieraea, 27: 255–286.

Martín, JL & Oromí P (1987) Tres nuevas especies hipogeas de Loboptera Brum. & W. (Blattaria, Blattellidae) y consideraciones sobre el medio subterráneo en Tenerife (Islas Canarias). Annls. Soc. Ent. Fr. (N. S.) 23(3): 315–326.

Mazzoni C, Ciofi C & Waterhouse R (2023). Biodiversity: An atlas of European reference genomes. Nature 619: 252–252. 10.1038/d41586-023-02229-w

